# Nuclear acetyl-CoA accumulation drives reversible RBP hyperacetylation and transcriptome-wide splicing shifts in age-equivalent Alzheimer’s disease neurons

**DOI:** 10.64898/2026.09.26.754222

**Authors:** Larissa Traxler, Oliver Borgogno, Kylie Champion, Joseph R. Herdy, Monika Dzieciatkowska, Julie A. Reisz, Ayk Petrosyan, Cat Siler, Colin C. Anderson, Elisa Gorostieta-Salas, Raffaella Lucciola, Douglas Galasko, Fred H. Gage, Angelo D’Alessandro, Jerome Mertens

## Abstract

Metabolic alterations and dysregulated alternative RNA splicing are central layers of Alzheimer’s disease (AD) pathogenesis, yet whether and to what extent these changes are mechanistically linked in adult human neurons in the aging brain remains unclear. To assess this, we generated induced neurons (iNs) by direct conversion of fibroblasts from sporadic AD patients and age-matched control donors. Fibroblast-derived iNs retaining donor-specific aging signatures have been shown to display several AD-related phenotypes. Metabolomic analysis revealed that AD iNs divert citrate toward ATP-citrate lyase (ACLY)-dependent citrate-to-acetyl-CoA flux, which drives increased nuclear levels of the high-energy metabolite acetyl-CoA. Acetyl-proteomics showed that acetyl-CoA accumulation leads to widespread hyperacetylation of RNA-binding proteins (RBPs) and splice regulators, suggestive of a direct mechanistic link between a metabolic shift and RNA processing. Indeed, single-cell long-read RNA sequencing identifies splicing alterations in AD-relevant transcripts whose isoform usage mirrors changes observed in post-mortem AD cortex samples. Subcellular acetyl-CoA imaging reveals that inhibition of ACLY enzymatic activity selectively reduces nuclear acetyl-CoA levels, and further improves citrate metabolic flux, normalizes RBP acetylation, and re-establishes a normal neuronal splicing landscape in AD iNs. Together, these findings reveal a nuclear acetyl-CoA-RNA processing axis that links age- and disease-related metabolic alterations to alternative splicing dysregulation in AD, and identify neuronal nuclear acetyl-CoA production as a causal and reversible node for restoring metabolic and RNA homeostasis in aging human AD neurons.

**Highlights:**

- Sporadic Alzheimer’s disease (AD) patient-derived directly induced neurons (iNs) shift metabolism toward nuclear acetyl-CoA production via ATP-citrate lyase (ACLY).
- Elevated nuclear acetyl-CoA causes hyperacetylation of RNA-binding proteins (RBPs).
- Patient iNs show pathogenic RNA splicing patterns similar to post-mortem AD brains.
- Inhibition of ACLY normalizes neuronal acetyl-CoA metabolism and restores RNA splicing homeostasis.

## Introduction

Key pathogenic drivers of Alzheimer’s disease (AD), including amyloid precursor protein (APP) and microtubule-associated protein tau (MAPT), are regulated at the level of alternative splicing, and genome-wide transcriptomic studies of the aging human brain are revealing a widespread shift in the neuronal splicing landscape in AD reaching beyond these canonical disease genes^1,2^. While splicing dysregulation is increasingly recognized as a fundamental feature of sporadic AD pathogenesis^1–7^, the upstream age-related cellular signals that trigger this global splicing shift remain unresolved. A coordinated change of this magnitude is unlikely to arise from a single dysfunctional splicing factor and instead points to an abundance-related change capable of acting on the entire RNA-binding proteome at once. Acetyl-CoA, a high-energy metabolic intermediate, links cellular energetic state to chromatin and proteome acetylation, thereby influencing transcription, RNA processing, and neuronal function^8–10^. Perturbations in acetyl-CoA homeostasis and protein acetylation have been previously associated with AD pathology^11–14^, but the cellular and subcellular mechanisms linking metabolic rewiring to AD-related RNA dysregulation remain unresolved.

Directly reprogrammed induced neurons (iNs) provide a uniquely powerful platform to dissect these mechanisms in a human, age-equivalent context. Unlike rejuvenated fetal-like iPSC-derived cell models, iNs retain donor-specific aging signatures, including epigenetic marks, transcriptional age, and mature neuronal isoform usage, in addition to the patients’ genetic makeup^15–18^. Importantly, adult donor-derived iNs endogenously express adult human 4R tau, develop a broad range of AD-relevant phenotypes, and recapitulate AD-associated transcriptional and splicing signatures observed in the human brain^15,17,19–24^. We previously showed that AD iNs undergo a Warburg-like metabolic shift toward glycolysis despite preserved mitochondrial function, raising the question of where the glycolysis-derived pyruvate is directed if not toward oxidative tricarboxylic acid (TCA) cycle metabolism^25^.

Here, using a cohort of directly converted iNs from sporadic AD patients and age-matched controls, we combine ¹³C_6_-glucose flux metabolomics, acetyl-proteomics, single-cell long-read RNA sequencing, and compartment-specific fluorescent acetyl-CoA sensor imaging to uncover that AD neurons divert citrate away from the TCA cycle and instead channel it toward ATP-citrate lyase (ACLY)-dependent production of nuclear acetyl-CoA. This subcellular redistribution and accumulation of nuclear acetyl-CoA drive widespread protein hyperacetylation, disproportionately affecting nuclear proteins, many of which orchestrate splicing. These acetylation changes cause a global shift in isoform splicing reminiscent of the splicing defects observed in post-mortem AD cortex. Pharmacological inhibition of ACLY normalizes mitochondrial citrate flux, nuclear acetyl-CoA levels, and RBP acetylation, and reverses AD-associated splicing changes.

## Results

### Generation of iNs from patient fibroblasts

We generated age-equivalent iNs by direct conversion of primary fibroblasts derived from a cohort comprised of 16 non-demented controls, 11 individuals with sporadic AD, and 2 familial AD cases carrying *APP-V717* and *PSEN1-A246E* mutations (Figure 1A, Figure S1A). Donor ages averaged 78 ± 7 years for controls, 79.5 ± 3 years for sporadic AD, and 55 ± 2.8 years for familial AD. Fibroblasts were directly converted to iNs using our established protocol^26^, which combines doxycycline-inducible overexpression of *ASCL1* and *NGN2* with a defined cocktail of small molecules that promotes neuronal fate and maturation. To remove intermediate cells, cultures were enriched for neuronal cells at three weeks following induction using fluorescence or magnetic activated cell sorting (FACS/MACS) based on cell-surface expression of PSA-NCAM^26^. Resulting iN cultures consist of a mixture of excitatory and inhibitory neurons with cortical identity, as previously defined^15,24^. Neuronal marker expression was robust across groups, with MAP2 expression detected in 95 ± 5% of control and 90 ± 9% of AD iNs, and Tau expression detected in 90 ± 7% of control and 87 ± 5% of AD iNs, with no significant differences between the groups (Figure 1B). We and others have previously reported that unlike rejuvenated iPSC-derived neuronal models, directly converted iNs retain key features of cellular aging, including epigenetic age, mitochondrial dysfunction, and transcriptional aging signatures, and adult brain-like mRNA splicing^16,18,22,23,27,28^. Notably, adult donor-derived iNs show adult *MAPT* splicing including Exon 10^15^, and show protein expression of both 3R and 4R Tau variants along their neurites (Figure 1B)^15,17^.

**Figure 1.**
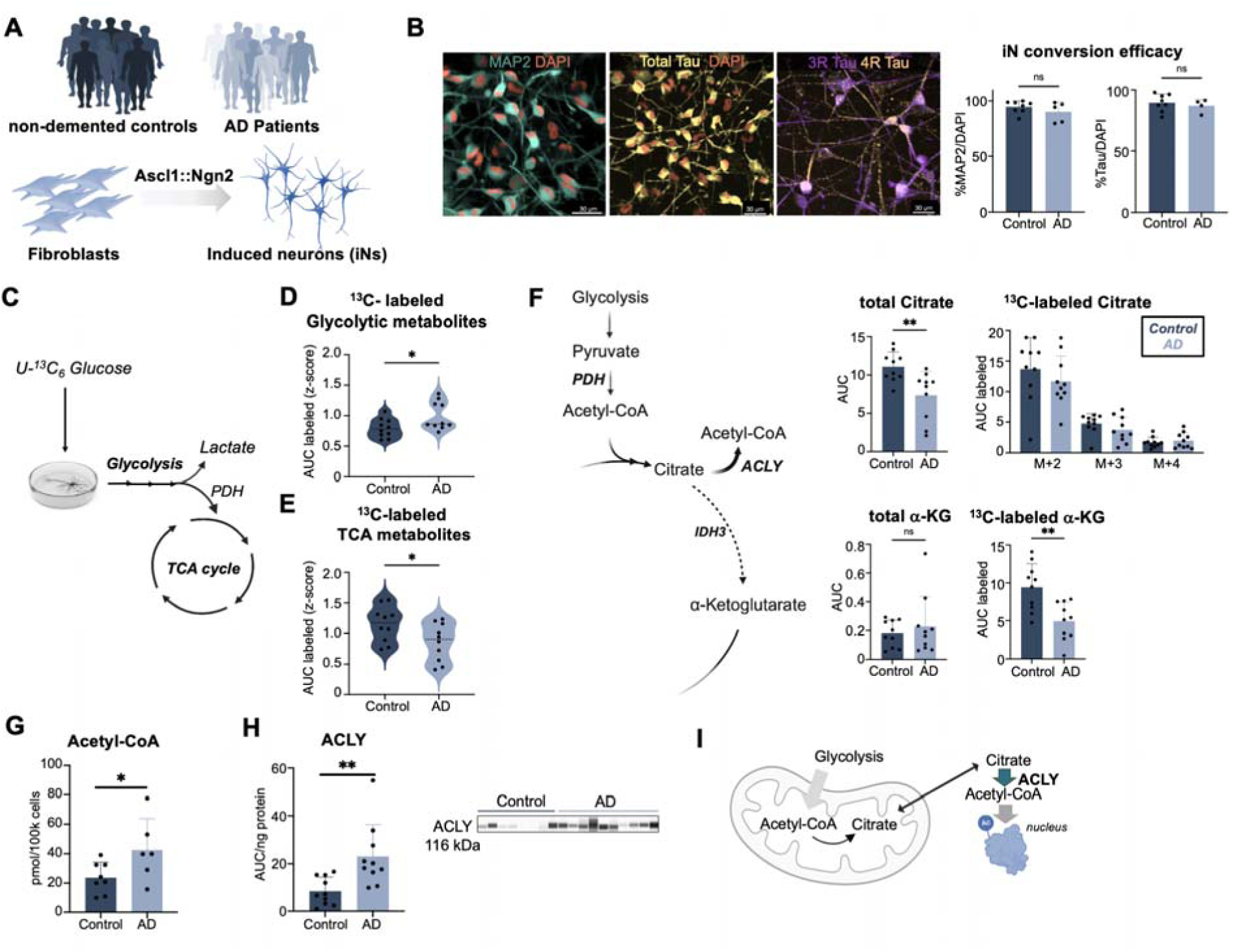
Metabolic re-wiring in AD iNs. (A) Schematic of the experimental approach. Fibroblasts were collected from a cohort of non-demented control donors and AD patients and directly converted into iNs by doxycycline-inducible overexpression of ASCL1 and NGN2. (B) Immunocytochemistry for MAP2, total Tau (Tau13), and 3R and 4R Tau isoforms. The percentage of cells expressing neuronal markers was quantified. Three images per donor (n=8 controls, n=4-5 AD) were analyzed, and each dot represents the average value per donor. Scalebar=30 µm. (C) Schematic of glucose flux tracing following 6 hours of incubation with ^13^C_6_-glucose. (D,E) Average z-score of isotopic labeling of glycolytic metabolites (D) and TCA cycle metabolites (E), quantified by UHPLC-MS. N=10 per group. (F) Total citrate and α-ketoglutarate (α-KG) levels, as well as the amount labeled following 6 hours of ^13^C_6_-glucose incubation quantified using UHPLC-MS metabolomics. N=10 per group. (G) Fluorescence-based assay for acetyl-CoA quantification in iNs, normalized to cell number. N=8 controls, n=6 AD. (H) ProteinSimple capillary western blot analysis of ACLY expression. N=10 controls, n=10 AD. (I) Proposed model of metabolic re-wiring in AD iNs. (B–H) Statistical significance was determined using unpaired two-tailed t-tests. Normality was assessed using Kolmogorov-Smironov and Shapiro-Wilk test. If normality was not met, a Mann Whitney test was performed. Each dot represents an independent donor. Data are presented as mean ± SD.

### Metabolic re-wiring causes a citrate-to-acetyl-CoA shunt in AD iNs

Aged AD neurons undergo a metabolic shift sharing similarities with the Warburg effect, characterized by increased aerobic glycolysis despite preserved mitochondrial function^25^. However, it remains unclear where mitochondrial pyruvate is directed toward supporting the TCA cycle. To interrogate this intrinsic metabolic rewiring, we performed ^13^C_6_-glucose flux tracing coupled to semi-quantitative UHPLC-MS metabolomics in purified iN cultures. Following fluorescent-activated cell sorting (FACS) and recovery, iNs were incubated for 6 hours with ^13^C_6_-Glucose to trace the flow of glucose-derived carbon through glycolysis and into the TCA cycle (Figure 1C). AD iNs exhibited increased incorporation of ^13^C into glycolytic intermediates, accompanied by a relative reduction in labeling of TCA cycle metabolites (Figures 1D-E). Interestingly, while global TCA cycle labeling was reduced, carbon flux into citrate was maintained in AD iNs (Figure 1F). Consistent with preserved mitochondrial entry of glycolytic carbon, AD iNs showed reduced inhibitory pyruvate dehydrogenase (PDH) protein phosphorylation (Figure S1B). Total citrate levels, however, were significantly decreased in AD, suggesting that citrate synthesis is maintained but its steady-state abundance is reduced due to downstream utilization (Figure 1F). Examination of downstream TCA cycle flux by quantifying labeling of additional metabolites further revealed that ^13^C incorporation into α-ketoglutarate (α-KG) was significantly reduced, demonstrating that indeed citrate production is uncoupled from downstream oxidative metabolism in AD iNs (Figure 1F). This pattern is consistent with a relative shift in isocitrate dehydrogenase (IDH) isoform usage, from IDH3 to IDH1, which alters carbon flux directionality between citrate and α-KG (Figure S1C). To better understand the discordance between maintained citrate production and reduced TCA progression, we assessed alternative citrate fates and found that AD iNs exhibited increased acetyl-CoA levels, which is consistent with enhanced citrate utilization outside the mitochondria (Figure 1G). In support of this, western blot analysis revealed increased protein levels of ACLY, the enzyme responsible for extra-mitochondrial conversion of citrate into acetyl-CoA (Figure 1H). These data indicate a metabolic rerouting of citrate from usage in the TCA cycle toward extra-mitochondrial acetyl-CoA production (Figure 1I).

### Hyperacetylation of RNA-binding proteins (RBPs) links metabolic re-wiring to aberrant splicing in AD iNs

Protein acetylation is largely controlled by local acetyl-CoA levels, as many lysine acetyltransferases operate near the limit of their affinity for this high-energy metabolite, making acetylation stoichiometry directly responsive to acetyl-CoA supply rather than solely to enzymatic activity^9^. Given the elevated acetyl-CoA levels observed in AD iNs as a likely direct consequence of the citrate shuttle, we next profiled the acetylated proteome by affinity enrichment of acetyl-lysine-containing peptides followed by mass spectrometry-based quantification. Partial least squares discriminant analysis (PLS-DA) on global acetylated protein abundance values clearly separated control from AD iNs (Figure S2A), characterized by a global protein hyperacetylation in AD iNs (Figure 2A). Differential analysis identified 57 significantly hyperacetylated and 23 significantly hypoacetylated proteins (p < 0.1) in AD iNs (Figure 2B). Mapping of the differentially acetylated proteins to the subcellular localization atlas of the Human Protein Atlas (proteinatlas.org^29^, 63 out of 80 differentially abundant proteins had an annotated localization) revealed that the largest fraction were nuclear proteins (34.9%), followed by cytosolic proteins (28.6%) and proteins that were both cytosolic and nuclear (11.1%) (Figure 2C). Gene ontology analysis (STRING GO-BP) showed a significant enrichment of processes related to mRNA processing and RNA splicing, while GO-CC terms highlighted spliceosomal and ribonucleoprotein complex associations (Figures 2D and S2B), and the POSTAR3 RNA-binding protein database^30^ confirmed ten hyperacetylated and four hypoacetylated RBPs (Figure 2B). To assess whether these changes reflected altered protein abundance, we performed quantitative global proteomics in four control and four AD iN samples, and unsupervised PCA revealed clear separation between the groups (Figure S2C). Differential protein abundance analysis (FDR < 0.1) identified 176 downregulated and 135 upregulated proteins (Figure S2D). Of the 80 proteins exhibiting significant changes in acetylation, only six (7.5%) also showed altered total protein abundance (Data S1). Thus, the vast majority of acetylation changes were not accompanied by corresponding changes in total protein abundance, indicating that acetylation changes are largely independent of protein abundance.

**Figure 2.**
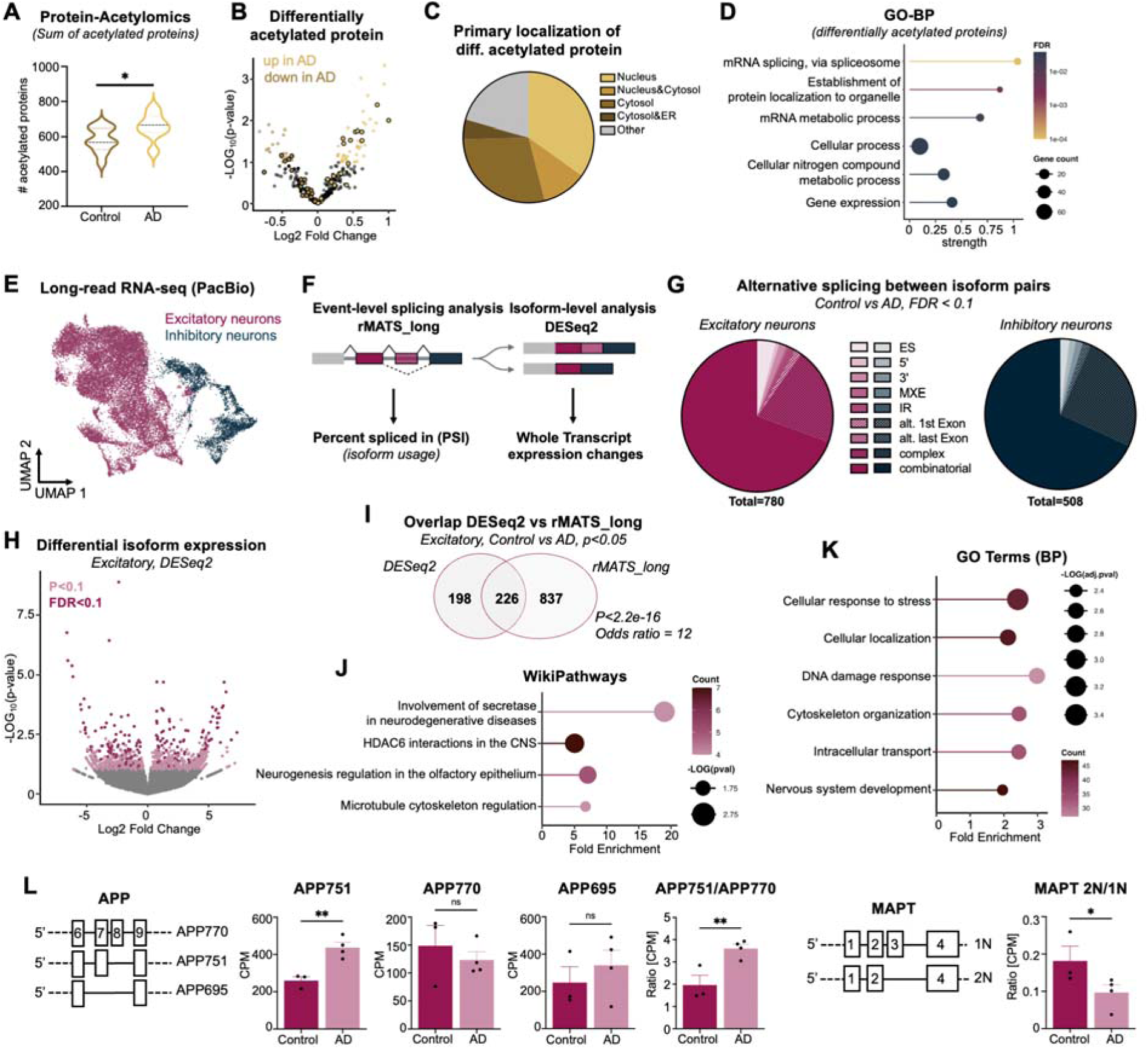
Hyperacetylation of RBPs and alternative splicing in AD iNs. (A) Acetylated proteins were enriched using the PTMScan Acetyl-Lysine Motif Kit and analyzed by mass spectrometry. Shown is the sum of acetylated proteins detected per condition. n = 11 control donors and n = 7 AD donors. Statistical significance was determined using an unpaired two-tailed t-test. (B) Differential acetylation analysis performed using limma on log-transformed data. (C) Primary annotated subcellular localization of differentially acetylated proteins based on Protein Atlas annotations. (D) Gene Ontology (GO) enrichment analysis performed using STRING. All significant terms with a false discovery rate (FDR) < 0.05 are shown. (E) UMAP visualization of single-cell long-read RNA sequencing (PacBio) data. Neuronal subtypes were annotated based on established marker genes. n = 3 control donors and n = 4 AD donors. (F) Schematic of the analysis workflow. (G) Alternative splicing events identified by rMATS-long in excitatory and inhibitory neurons following pseudobulk aggregation of each cell type. (H) DESeq2 differential transcript expression analysis showing all detected transcripts; transcripts with p < 0.1 are highlighted in light pink and transcripts with FDR < 0.1 in dark pink. (I) Overlap of significantly altered transcripts identified by DESeq2 and rMATS-long analysis. Statistical significance was calculated using Fisher’s exact test. (J,K) DAVID functional annotation analysis of overlapping genes that are alternatively spliced in AD iNs identified by DESeq2 and rMATS-long, showing the top four enriched WikiPathways terms (J) and top six enriched Gene Ontology Biological Process terms (K). All significant terms are shown in Figure S2H. (L) Representative genes differentially spliced in AD iNs. Statistical significance was determined using unpaired two-tailed t-tests. Normality was assessed using Kolmogorov-Smironov and Shapiro-Wilk test. Each dot represents an independent donor. Data are presented as mean ± SD.

### Metabolic re-wiring is associated with widespread isoform dysregulation across excitatory and inhibitory AD iNs

To determine whether metabolic and post-translational changes translate into a global shift in RNA processing, we performed pooled single-cell long-read RNA sequencing (PacBio) of three control and four AD iN samples. Following magnetic-activated cell sorting (MACS)-based neuronal enrichment and replating to allow recovery, cells were harvested and processed using the 10x Genomics single-cell workflow. All seven donor samples were pooled into a single 10x run, and PacBio libraries were generated from the resulting 3’-barcoded cDNA, enabling full-length transcript sequencing. Following long-read sequencing, cells were computationally demultiplexed based on donor-specific single-nucleotide polymorphisms (SNPs), yielding an average of 1,630 ± 307 high-quality cells per donor, with an average read length across donors of 1,009 ± 566 bp (maximum 7,505 bp). Similar to previously reported numbers based on imaging and short-read single cell RNAseq, our iN cultures consist predominantly of excitatory neurons (79.5% ± 6.4%) and a smaller fraction of inhibitory neurons (20.5% ± 6.4%) (Figure 2E), which were computationally separated before downstream RNA isoform analysis^15,21^. To assess alternative splicing in both neuronal subtypes separately, we aggregated reads by cell identity and applied two complementary analytical approaches: (i) Event-level differential splicing using rMATS_long leveraging transcript discovery using ESPRESSO, and (ii) Isoform-level differential expression using DESeq2 (Figure 2F). The rMATS_long models isoform usage as a beta-binomial distribution of relative read proportions, making it particularly well suited for detecting differential splicing because it directly tests whether the relative contribution of alternative isoforms from the same gene shifts between conditions while accounting for biological and technical variability^31,32^. This compositional framework enables detection of changes in transcript processing that are distinct from changes in overall gene expression. However, because isoform usage is modeled as a relative quantity, it can obscure coordinated changes in the absolute abundance of individual isoforms, motivating complementary isoform-resolved differential expression analysis. In excitatory neurons, rMATS_long identified 780 significant differential splicing events (FDR < 0.1), whereas inhibitory neurons showed 508 events (Figure 2G). At the isoform level, DESeq2 identified 199 and 51 differentially expressed isoforms in excitatory and inhibitory neurons, respectively (FDR < 0.1) (Figures 2H, S2E, S2G, and S2I). Notably, there was minimal overlap in the genes exhibiting differential splicing or isoform expression between excitatory and inhibitory neurons, suggesting largely distinct, neuronal subtype-specific transcriptomic alterations (Figure S2F). For pathway-level interpretation, we included transcripts with nominal significance (p < 0.05) and focused on overlapping hits between both methods to increase stringency (Figures 2I and S2I). Pathway enrichment analysis of excitatory neuron-specific isoform changes using both WikiPathways and GO-BP revealed enrichment in pathways related to secretase in neurodegenerative diseases, central nervous system (CNS) acetylation, neurogenesis, microtubule cytoskeletal regulation, stress response, intracellular transport, and localization pathways (Figures 2J-K and S2H), with five key genes overlapping between WikiPathways and GO-BP results including the AD-relevant genes APP, MAPT, and RAC1 (Figures 2L, S2K)^2,33^. Consistent with prior short-read transcriptomic analyses of post-mortem brain tissue^2^, AD iNs exhibited increased expression of the APP751 isoform and a shift from APP770 to APP751 (Figure 2L). Similarly, MAPT isoform analysis revealed a shift from 2N to 1N tau isoforms (Figure 2L), and RAC1 showed reduced expression of the RAC1B isoform (Figure S2K). Additionally, AD iNs showed higher expression of ABL1a and the truncated isoform of MAT1B (Figure S2K). In contrast, inhibitory neurons did not show significant enrichment in WikiPathways, and GO-BP analysis pointed towards altered RNA splicing programs (Figure S2J). Inhibitory neurons also exhibited isoform changes consistent with post-mortem datasets, including alterations in KIDINS220 splicing (Figure S2L), and differential isoform usage of multiple RNA-processing genes, including RBFOX3, where AD iNs increased expression of an isoform lacking exon 15, previously implicated in altered nuclear localization (Figure S2L)^34,35^.

### Inhibition of ACLY restores subcellular acetyl-CoA homeostasis and mitochondrial metabolism in AD iNs

Our findings suggested that metabolic rewiring toward increased extra-mitochondrial acetyl-CoA production contributes to RBP hyperacetylation and splicing dysregulation in AD iNs. To establish whether elevated total acetyl-CoA levels reflect global or compartment-specific alterations, we sought to resolve subcellular acetyl-CoA dynamics using internally controlled PancACe acetyl-CoA sensors targeted to the cytoplasm, mitochondria, or nucleus (Figure 3A)^36^. PancACe sensors were co-transduced with synapsin-driven dsRed and, in line with the acetyl-proteome data, revealed a selective accumulation of nuclear acetyl-CoA in AD iNs, whereas cytosolic acetyl-CoA remained unchanged and mitochondrial acetyl-CoA showed a trend toward reduction (Figure 3B). To identify metabolic interventions that specifically reduce nuclear acetyl-CoA while preserving mitochondrial pools, we tested inhibition of ACLY using SB204990 (ACLYi) and of acyl-CoA synthetase short chain family member 2 (ACSS2) using Ac-CoA Synthase Inhibitor 1 (ACSS2i), modulators of PDH activity using dichloroacetate (DCA), and activators of sirtuin-dependent deacetylation pathways using SRT1720 supplemented with nicotinamide riboside (SRT1720+NR) (Figure S3A)^36^. None of the treatments significantly altered cytosolic acetyl-CoA levels, but ACLYi, DCA, and SRT1720+NR reduced nuclear acetyl-CoA levels (Figure S3A). Notably however, only ACLYi preserved mitochondrial acetyl-CoA, whereas SRT1720+NR caused a pronounced reduction and DCA showed a trend toward decreased mitochondrial acetyl-CoA (Figures S3A-B). Prolonged ACLYi treatment up to 10 days did not adversely affect neuronal morphology or identity, as MAP2 and Tau expression remained unchanged following treatment (Figure 3C). UHPLC-MS analysis further demonstrated that ACLYi normalized elevated acetyl-CoA levels without globally suppressing acetyl-CoA production in AD iNs, while it did not significantly alter acetyl-CoA levels in control iNs (Figure 3D), or affecting other CoA intermediates (Figures S3C-D). When applied to AD patient iNs, PancACe imaging confirmed that ACLYi effectively reduced pathological nuclear acetyl-CoA accumulation (Figure 3E), while cytosolic acetyl-CoA remained unchanged across conditions (Figure S3E), and mitochondrial acetyl-CoA levels even increased following ACLY inhibition (Figure 3E). This is consistent with reduced citrate export from mitochondria and enhanced substrate availability for the TCA cycle. To determine whether these metabolic changes translated into improved TCA activity, we quantified citrate abundance and TCA cycle flux. AD iNs exhibited significantly reduced citrate levels, which were restored following ACLYi treatment (Figures 1F and 3F). Consistently, ^13^C_6_-glucose tracing revealed increased labeling of α-KG upon ACLYi treatment, indicating enhanced TCA cycle activity (Figure 3G). Together, these findings demonstrate that ACLY inhibition restores subcellular acetyl-CoA balance and improves TCA cycle flux in AD iNs (Figure 3H).

**Figure 3.**
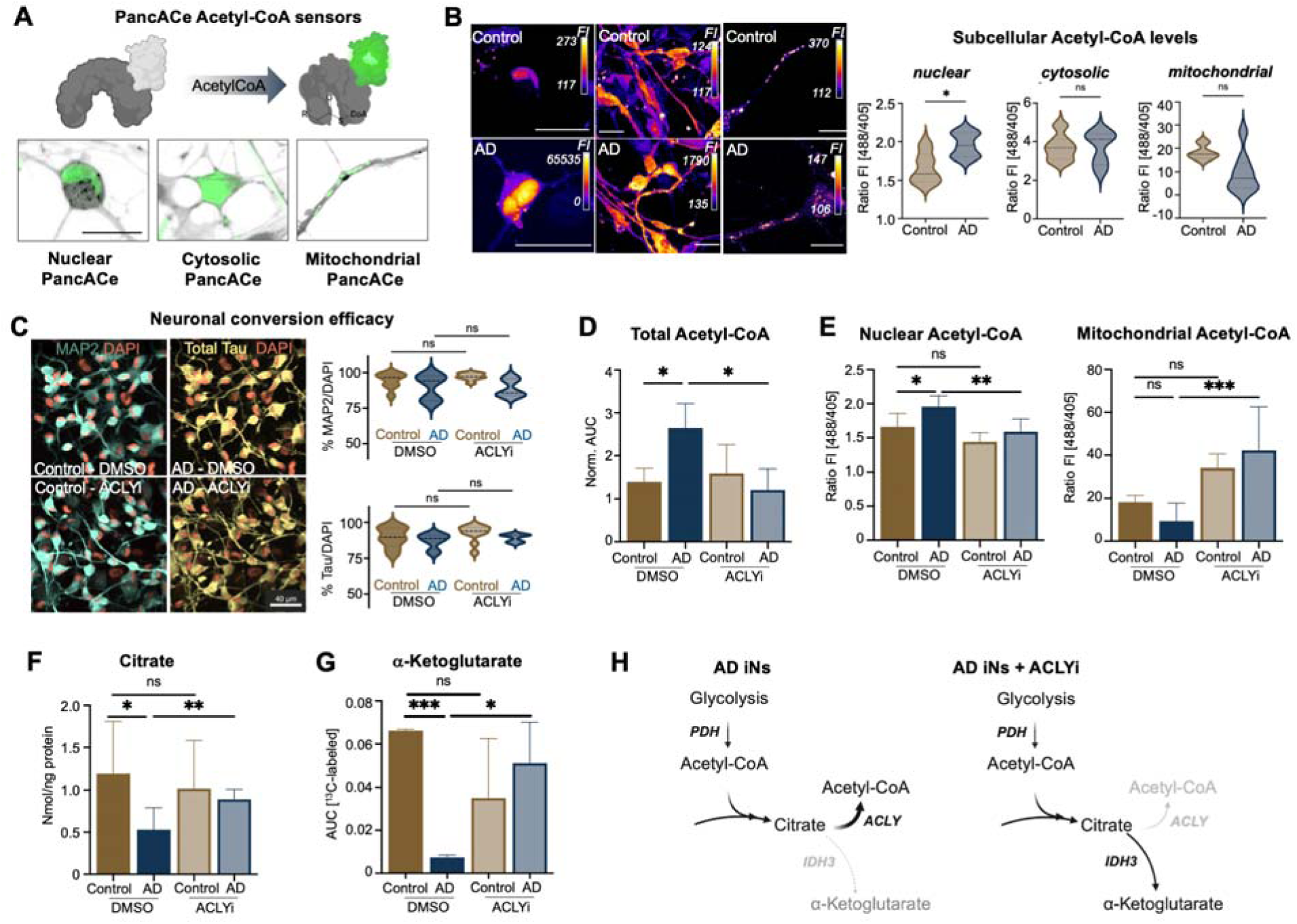
Inhibition of ACLY restores subcellular acetyl-CoA homeostasis and mitochondrial metabolism in AD iNs. (A) Schematic and representative live-cell images of compartment-specific PancACe acetyl-CoA sensors. iNs were co-transduced with synapsin::dsRed (grey) and one of the PancACe sensors (green). Scale bar = 20 µm. (B) Representative images and quantification of live-cell imaging of nuclear, cytosolic, and mitochondrial PancACe sensors in control and AD iNs. Sensor intensity was measured using 488 nm excitation and normalized to the internal control signal excited at 405 nm. nuclear PancACe: n = 9 control donors, n = 5 AD donors; Scale bar = 25 µm. cytosolic PancACe: n = 12 control donors, n = 5 AD donors; mitochondrial PancACe: n = 4 control donors, n = 8 AD donors. Scale bar = 15 µm. Statistical significance was determined using unpaired two-tailed t-tests. (C) Effect of ACLY inhibition (10 µM SB-204990) on neuronal marker expression. iNs were treated with either DMSO or ACLYi for 10 days prior to analysis and stained for MAP2 and Tau. Statistical significance was determined using ordinary one-way ANOVA followed by Sidak’s multiple comparisons test. Scale bar = 40 µm. n = 8 control donors and n = 5 AD donors. (D) UHPLC-MS quantification of acetyl-CoA levels at baseline (DMSO) or following ACLYi treatment (10 µM, 7 days). n = 5 control donors and n = 3 AD donors. Statistical significance was determined using ordinary one-way ANOVA (F(3,12) = 4.8) followed by Sidak’s multiple comparisons test. (E) Quantification of PancACe sensors at baseline and after ACLYi. ordinary one-way ANOVA followed by Sidak’s multiple comparisons test. Nuclear PancACe: F(3,24) = 8.3; n = 9 control DMSO, n = 5 AD DMSO, n = 6 control ACLYi, and n = 8 AD ACLYi donors. Mitochondrial PanACe: F(3,21) = 9.6; n = 4 control DMSO, n = 8 AD DMSO, n = 5 control ACLYi, and n = 8 AD ACLYi donors. (F) Fluorescence-based citrate assay. Citrate concentrations were calculated from a standard curve and normalized to protein content determined by BCA assay. n = 8 control donors (DMSO/ACLYi) and n = 6 AD donors (DMSO/ACLYi). Statistical significance was determined using ordinary one-way ANOVA (F(3,24) = 2.4) followed by Sidak’s multiple comparisons test. A paired two-tailed t-test was additionally performed to compare AD DMSO versus AD ACLYi conditions. (G) UHPLC-MS glucose flux tracing following 6 hours of incubation with ^13^C_6_-glucose. Cells were treated with either DMSO or ACLYi (10 µM) for 7 days prior to analysis. n = 2 per group. Statistical significance was determined using unpaired two-tailed t-tests. (H) Proposed model of the effect of ACLY inhibition on metabolic re-wiring in AD iNs. Data are presented as mean ± SD.

### ACLY inhibition reverses hyperacetylation of RBPs and restores a healthy RNA isoform landscape

To determine whether normalizing extra-mitochondrial acetyl-CoA in AD iNs affects downstream phenotypes, we repeated acetyl-proteomics in healthy control and AD iNs under baseline conditions (DMSO) and following ACLYi treatment. In this independent experiment, and consistent with the previous experiment, DMSO-treated AD iNs displayed global protein hyperacetylation, and ACLYi robustly reversed this phenotype across all donors to levels similar to healthy iNs (Figure 4A). Treatment of iNs with a broad deacetylase inhibitor cocktail targeting class I/II/III histone deacetylases and SIRT (Trichostatin A, EX-527, Nicotinamide, and Sodium butyrate) led to a global increase in protein acetylation, confirming that these changes reflect true biological modulation of protein acetylation (Figure S4A). Normalization to total protein using the non-binding fraction after acetyl pulldown yielded consistent results (Figure S4B), indicating that the observed changes are not driven by differences in total protein abundance. Differential acetylation analysis confirmed a pronounced hyperacetylation phenotype in AD iNs, with 528 proteins significantly hyperacetylated compared to controls (FDR < 0.1; Figure 4B). Upon ACLY inhibition, 211 proteins showed significantly reduced acetylation (FDR < 0.1), and after 7 days of treatment, the acetyl-proteomic profiles of AD iNs were indistinguishable from control iNs, with no remaining significantly hyper- or hypoacetylated proteins (Figure 4B). Intersection of our dataset with the POSTAR3 database confirmed the initial results in that RBPs were prominently hyperacetylated in AD iNs. Out of 111 detected RBPs, 26 were significantly hyperacetylated (FDR < 0.1) and 51 at nominal significance (p < 0.01) (Figure 4C). ACLYi markedly reversed RBP hyperacetylation in AD iNs, with 10 RBPs significantly reduced (FDR < 0.1) and 24 at nominal significance (p < 0.1); all reversing the AD phenotype in directionality (Figure 4C). ACLYi had minimal effects on control iNs RBP acetylation. These findings indicate that ACLY activity directly modulates RBP acetylation in the disease context.

**Figure 4.**
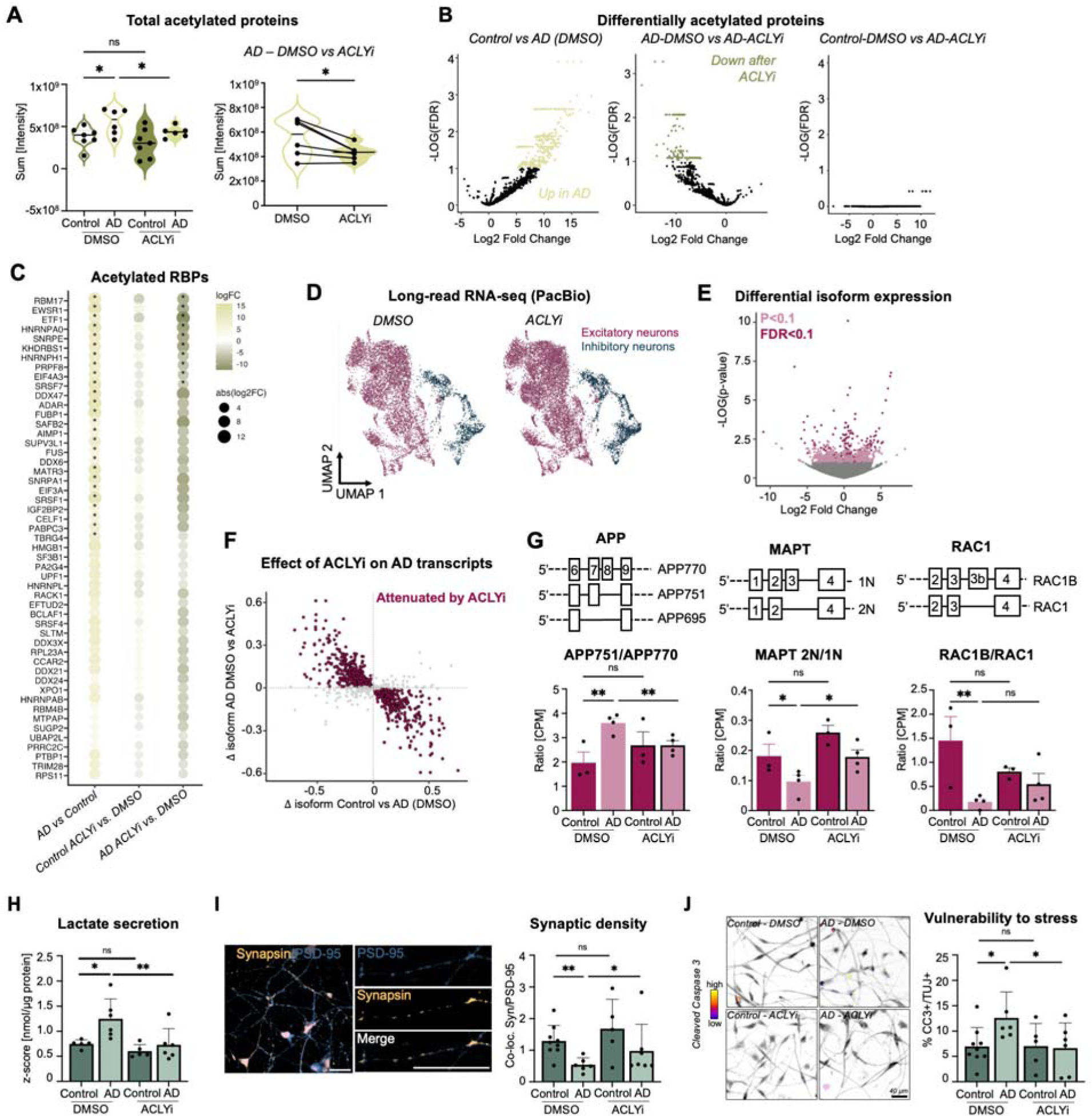
ACLY inhibition reverses hyperacetylation of RBPs and restores a healthy RNA isoform landscape. (A) Acetylated proteins were enriched using the PTMScan Acetyl-Lysine Motif Kit and analyzed by mass spectrometry. iNs were MACS-purified and treated with either DMSO or ACLYi for 7 days. Shown is the sum of intensities of acetylated proteins detected per condition. n = 7 control donors (DMSO/ACLYi) and n = 6 AD donors (DMSO/ACLYi). Statistical significance was determined using ordinary one-way ANOVA (F(3,22) = 4.0). Paired two-tailed t-tests were additionally performed to compare matched AD DMSO and AD ACLYi samples, indicated by connecting lines. (B) Differential acetylation analysis performed using limma on log-transformed data comparing control versus AD under DMSO conditions, AD DMSO versus AD ACLYi, and control DMSO versus AD ACLYi conditions. (C) RNA-binding proteins (RBPs) were filtered based on the POSTAR3 database and ranked by significance in the AD versus control DMSO comparison. All RBPs with p < 0.01 are shown; RBPs with FDR < 0.1 are indicated by asterisks. Dot color and size represent log_2_ fold change. (D) UMAP visualization of single-cell long-read RNA sequencing (PacBio) data from iNs treated with DMSO or ACLYi. Neuronal subtypes were annotated based on established marker genes. n = 3 control and n = 4 AD donors. (E) DESeq2 differential transcript expression analysis in excitatory neurons comparing AD DMSO versus AD ACLYi conditions. All detected transcripts are shown; genes with p < 0.1 are highlighted in light pink and genes with FDR < 0.1 in dark pink. (F) Correlation of isoform changes between control versus AD (DMSO; x-axis) and AD DMSO versus AD ACLYi (y-axis). Each dot represents a transcript significantly altered (p < 0.05) in the control versus AD DMSO comparison. Red dots indicate transcripts shifted in the opposite direction following ACLYi treatment (rescue). (G) Representative genes differentially spliced in AD iNs. Statistical significance was determined using ordinary one-way ANOVA followed by paired two-tailed t-tests comparing matched AD DMSO and AD ACLYi samples. APP: F(3,10) = 4.1. MAPT: F(3,10) = 6.3. RAC1: F(3,10) = 4.4. Each dot represents an independent donor. Data are presented as mean ± SD. (H) Lactate was measured in the supernatant using fluorescence quantification kit. n=5 control and n=6 AD donors. Statistical significance was determined using ordinary one-way ANOVA followed by paired two-tailed t-tests comparing matched AD DMSO and AD ACLYi samples. F(3,18) = 5.9. (I) Representative immunostaining and quantification of synapsin and PSD-95. Co-localization of the pre- and post-synaptic marker were used to determine synaptic density. n=9 control and n=7 AD donors. Statistical significance was determined using ordinary one-way ANOVA followed by paired two-tailed t-tests comparing matched AD DMSO and AD ACLYi samples. F(3,23) = 3.1. Scalebar = 20 µm. (J) Representative immunostaining and quantification of cleaved caspase 3 and β-III-tubulin (TUJ) in iNs treated with ABT-737 (0.15 µM) for 6 hours before fixation. n=9 control-DMSO, n= 6 AD-DMSO and AD-ACLYi, n=5 control-ACLYi. Statistical significance was determined using ordinary one-way ANOVA followed by paired two-tailed t-tests comparing matched AD DMSO and AD ACLYi samples. F(3,22) = 2.5. Scalebar = 40 µm.

We next performed single-cell long-read RNA sequencing following ACLY inhibition (Figure 4D) to test whether these changes translate into restoration of the RNA isoform landscape. In excitatory neurons, ACLYi significantly altered 133 transcripts (FDR < 0.1) and 422 at nominal significance (p < 0.05) (Figure 4E), whereas effects in inhibitory neurons were more modest (16 transcripts at FDR < 0.1; Figure S4C), likely reflecting reduced statistical power due to lower cell numbers and exclusion of one donor. To assess disease-specificity, we tested whether ACLYi preferentially rescues AD-associated isoform changes. Disease-associated transcripts (p < 0.05, Control vs AD) were ∼2-fold more likely to be rescued by ACLYi in excitatory neurons (rMATS_long: OR = 2.35, p = 3.7×10^⁻48^; DESeq2: OR = 1.71, p = 1.1×10^⁻8^; Figures 4F and S4D), and in inhibitory neurons (rMATS_long: OR = 1.76, p = 8.6×10^⁻20^; DESeq2: OR = 2.55, p = 2.0×10^⁻7^; Figure S4E). This enrichment was robust across a range of effect size thresholds. Importantly, ACLY inhibition corrected key disease-relevant isoform changes identified earlier: In excitatory neurons, it reduced expression of APP751/APP770 to baseline levels, restored the MAPT 2N/1N ratio, and showed a trend toward normalization of RAC1B/RAC1, ABL1a/b and MAP1B (Figures 4G, S4F). In inhibitory neurons, ACLY inhibition rescued RBFOX isoform expression, although no beneficial effect on KIDINS220 was observed (Figure S4G).

To probe to what extent these molecular changes translate into functional rescue, we tested the established AD iN dedifferentiation phenotypes characterized by increased glycolysis, reduced synaptic density^15,25^, and heightened stress vulnerability^15,23,25^. Indeed, ACLY inhibition reversed these features, normalizing reducing anaerobic glycolysis indicated by reduced lactate secretion, increasing synaptic density, and reducing stress-induced cell death (Figures 4H-4J), which is consistent with restoration of a healthy neuronal state.

## Discussion

A key strength of this study is the use of directly converted iNs, which retain the age-associated molecular and metabolic features of the donors they came from. Unlike rejuvenated iPSC-derived neurons, which erase donor age and adopt an embryonic identity during reprogramming, iNs allow for the study of human-specific, age-dependent, and disease-related process directly in adult, AD patient-derived neurons^15,17,19,21,25,27,37^. Splicing undergoes extensive developmental reprogramming, with MAPT, APP, and many other transcripts exhibiting distinct isoform profiles in fetal versus adult brain. The acquisition of adult-enriched isoforms, such as 4R-MAPT and APP695, is a defining feature of adult neurons^38,39^. Importantly, iNs endogenously express these adult splicing programs, which become a prerequisite when studying AD-associated RNA splicing changes and comparing them directly to AD post-mortem brain data. The finding that the resulting splicing changes recapitulate key changes reported in post-mortem AD cortex confirm the notion that iNs capture age-dependent AD disease biology related to RNA splicing.

We previously showed that donor aging, at the absence of pathology, impairs mitochondrial function in iNs^27^, and that AD patient-derived iNs undergo a Warburg-like shift toward anaerobic glycolysis compared to age-matched controls, despite maintained mitochondrial function^25,40^. Our data now reveal that mitochondrial pyruvate entry through PDH remains active and citrate ^13^C-labeling is preserved, but that downstream citrate flux is redirected, with a greater fraction being exported from mitochondria and converted to extra-mitochondrial acetyl-CoA through elevated ACLY activity. This is also consistent with the accompanying IDH3-to-IDH1 isoform shift, which can favor reductive carbon flux from α-KG toward citrate^41,42^. This subcellular imbalance of acetyl-CoA is distinct from a general bioenergetic failure^25^, but instead appears because of a redistribution program where changes in metabolic flux lead to increased nuclear acetyl-CoA levels and elevated protein-acetylation in the nucleus.

Alternative splicing dysregulation is increasingly recognized as a widespread and functionally consequential feature of AD. Multiple post-mortem transcriptomic studies have documented extensive isoform alterations across cortical regions, far exceeding the scope attributable to any single splicing regulator ^2,4,43–49^. While prior work has focused heavily on specific splice factors, most prominently TDP-43, the magnitude and diversity of splicing changes observed in AD brain strongly suggests that many splicing factors become dysfunctional in the disease context^1–7^. Our findings provide a mechanistic explanation for this splicing diversity by linking metabolic rewiring to widespread RBP hyperacetylation and consequent disruption of RNA processing. Rather than a single factor going rogue, our data point to mechanism where the over-abundance of a high-energy metabolite drives a shift in a large fraction of the nuclear RBP proteome at once. Acetylation of RBPs can disrupt their RNA recognition and binding^50–55^, localization^51,56^, and stability^57^. Notable, many RBPs are highly dosage-sensitive, where both overexpression and knock-down is toxic to neurons, suggesting that their function is tightly controlled a the post-translational level^58^. Importantly, acetylation has also been implicated as an early pathogenic event preceding disease-associated phosphorylation and aggregation of proteins such as TDP-43 and Tau across neurodegenerative diseases^12,59,60^. Together, these observations suggest that the extensive splicing abnormalities observed in AD neurons and post-mortem brain may arise from the combined dysfunction of multiple RBPs, each perturbed by a shift towards aberrant hyperacetylation in distinct but convergent ways.

ACLY has previously been proposed as a driver of pathogenic protein acetylation in neurodegeneration^61^. Here, we show that pharmacological ACLY inhibition selectively reduces the elevated nuclear acetyl-CoA pool in AD iNs without disrupting global acetyl-CoA metabolism, thereby restoring mitochondrial citrate utilization. ACLY inhibition further exerts broad effects on the protein acetylome, normalizing acetylation patterns toward healthy neuronal states. Strikingly, this metabolic intervention rescues AD-associated isoform usage with high specificity, as transcripts dysregulated in AD reverted toward control splicing patterns following ACLY inhibition, while most other AD-unrelated transcripts remained unaffected. This directional rescue provides causal evidence that altered acetyl-CoA metabolism directly orchestrates splicing dysfunction in AD neurons. Importantly, these effects occur without compromising neuronal viability, suggesting that modulation of subcellular acetyl-CoA pools may represent a therapeutic strategy to restore both RNA and metabolic homeostasis in neurodegeneration. Together, our findings support a model in which AD neurons exist in a maladaptive metabolic state characterized by elevated nuclear acetyl-CoA, widespread RBP hyperacetylation, and destabilization of neuronal splicing programs.

## Limitations of the study

Studying AD pathological mechanisms in isolated neurons lets us resolve cell-autonomous AD mechanisms that could otherwise be masked by glial contributions in a more complex system. However, neuronal monocultures lack the broader brain context in which glia could either buffer these metabolic shifts or amplify the resulting pathology, and our model system cannot resolve between these possibilities. Nevertheless, direct metabolic profiling in post-mortem human brain remains difficult given the lability of metabolites like acetyl-CoA and confounding shifts in cellular composition across disease stages, underscoring the value of a living, cell-type-resolved model such as iNs for capturing important dynamic regulatory mechanisms. This study is based on a well-characterized, clinically stratified cohort in which metabolic effects were consistent across donors; however, AD is now understood to be etiologically heterogeneous rather than a single disease process, and larger cohorts with targeted assays and biomarkers will be needed to capture the full extent of that variability. While our findings establish a strong association between RBP hyperacetylation and splicing dysfunction, the precise mechanisms by which acetylation alters individual RBP function, including RNA binding, protein interactions, localization, and phase behavior, remain to be resolved. Identifying which RBPs sit at key mechanistic splicing nodes will be an important next step, both as additional therapeutic targets and as biomarkers, to facilitate moving this concept closer to therapeutic application in neurodegenerative disease.

## Acknowledgements

We want to thank Anna McTigue, and Mario Medina for technical support in this project. We would like to thank the Stem Cell Genomics Core at the Sanford Stem Cell Institute for providing sequencing services. We would further like to thank the Stem Cell Core Facility (RRID:SCR_014850), the Flow Cytometry Core Facility (RRID:SCR_014839), and the Mass Spectrometry Core (RRID:SCR_014843) at the Salk Institute for Biological Studies. This work was supported by the National Institute on Aging awards R01-AG085634, RF1-AG056306, RF1-AG095193, P30-AG062429, P30CA014195, S10-OD023689 and U19-AG023122, the California Institute for Regenerative Medicine (CIRM) awards DISC0-17507 and DISC4-19291, the BrightFocus Foundation awards A2019562S and A2022024F; the Live like Lou Foundation, the Theodor Koerner Fonds, L’Oreal Austria/OeUK/OeAW Stipend, the Freedom Together Foundation, the Dolby Foundation, and the American Federation for Aging Research (AFAR) award 30376715.

## Author contributions

L.T. and J.M. designed the study, L.T., K.C., A.P., and E.G.S. conducted tissue culture work, L.T., D.S., C.C.A., J.A.R. and A.D. performed metabolomics analyses, L.T., M.D., E.G.S., and F.H.G. designed proteomics experiments and performed proteomics analyses, K.C. and C.S. performed molecular biology analyses, L.T., J.R.H., R.L. and O.B. performed bioinformatic analyses and visualized data, D.G. performed clinical characterization of the cohort and obtained skin punch biopsies, L.T. and J.M. wrote the paper. All authors have thoroughly discussed this project and helped improve the paper.

## Declaration of interests

The authors declare no competing interests.

### Supplemental information

Document S1. Figures S1-S4

Data S1. Significant proteins

## STAR METHODS

### RESOURCE AVAILABILITY

#### Lead contact

Further information and requests for resources and reagents should be directed to and will be fulfilled by the lead contact Jerome Mertens

#### Materials availability

This study did not generate new unique reagents.

#### Data and code availability

All rawdata will be uploaded to ArrayExpress and Metabolomics Workbench.

## EXPERIMENTAL MODEL AND SUBJECT DETAILS

### Fibroblasts and iNs

Human fibroblasts were obtained from the UCSD Shiley-Marcos Alzheimer’s disease research center (ADRC) and provided written informed consent. All procedures were approved by the local human subjects committee. The cohort consisted of 16 control donors aged 65-87 years, of whom 50% were female, and 13 AD patients aged 53-85 years, of whom 46% were female. The cohort was genetically balanced with respect to APOE genotype, including APOE2/3 (7%), APOE3/3 (41%), APOE3/4 (41%), and APOE4/4 (10%) (Figure S1A). Fibroblasts were cultured in DMEM containing 15% fetal bovine serum and 0.1% non-essential amino acids at 37 °C with 5 % CO_2_. They were transduced with a lentivirus pLVXUbC-rtTA-Ngn2:2A:Ascl1 (Addgene: #127289) and selected with puromycin (1 µg/ml) as previously described^26^. To initiate conversion, puromycin-selected fibroblasts were pooled 3:1 and, after 24 hours, medium was changed to neural conversion medium for three weeks. Neural conversion medium is based on DMEM:F12 and Neurobasal (1:1), supplemented with N2 supplement (1x, ThermoFisher), B27 supplement (1x, ThermoFisher), doxycycline (2 µg/ml, Sigma Aldrich), Laminin (1 µg/ml, Sigma Aldrich), dibutyryl-cyclic-AMP (100 µg/ml, Santa Cruz), DMH1 (5 µM, Biotechne, Medchem Express), LDN-193189 (0.5 µM, Sanova Pharma), A83-1 (0.5 µM, Santa Cruz), CHIR99021 (3 µM, LC Laboratories), forskolin (5 µM, LC Laboratories) and SB-431542 (10 µM, MedChem). After three weeks of conversion, cells were detached with TrypLe (ThermoFisher) and either plated on Geltrex-coated (ThermoFisher) µ-slides (ibidi) or FACS/MACS sorted. For sorting, cells were detached with TrypLE and stained with PSA-NCAM-PE or PSA-NCAM-microbeads (Milteny Biotec) in PBS containing 5 % FBS. Cells were FACS sorted in PBS containing Rock-inhibitor (10 µM) and DAPI, or MACS sorted in either PBS or neuronal conversion medium supplemented with Rock-inhibitor and z-VAD(Ome)-FMK. Sorted cells were plated in conversion media containing Rock-inhibitor and z-VAD(OMe)-FMK. Cells were treated with ACLY inhibitor SB204990 (10 µM, MedchemExpress) for 1 week after sorting until harvest.

## METHOD DETAILS

### Immunocytochemistry

Cells were fixed with 4 % paraformaldehyde and stained in PBS containing 0.05 % Triton-X100 and 5 % FBS. Cells were incubated with primary antibodies NeuN (CellSignaling, 1:200, #24307T), MAP2 (Abcam, ab5392), β-III-tubulin (Novus Biologicals, NB100-1612), anti-tau 3-repeat isoform RD3 (EMD Milipore, 1:300, #05-803), anti-4R tau antibody (Abcam, 1:300, ab218314, anti-tau), and anti-tau (Tau-13, BioLegend, 1:500, #835201) at 4°C overnight and incubated with secondary antibodies for two hours at room temperature, followed by 10 minutes of DNA staining with DAPI (ThermoFisher, 300nM, D21490). Images were taken with the ECHO Revolve Microscope and the Andor Dragonfly and analyzed in FIJI.

### Mass spectrometry metabolomics

A total of 150,000 iNs were FACS/MACS purified and re-plated in BrainPhys media supplemented with small molecules as described in the neuronal conversion media. After a 48 hours recovery period, iNs were supplemented with 2.5 mM or 1.3 mM ^13^C_6_-Glucose (Santa Cruz, sc-239643A; Millipore Sigma, #389374). To quantify labeling of glycolysis and TCA cycle metabolites, iNs were incubated for six hours with heavy-labeled metabolites, to quantify labeling of CoA-metabolites iNs were incubated for 16 hours before collecting cell pellets. Cell pellets were resuspended in Lysis Buffer (50:30:20 MeOH:Acetonitrile(ACN):H2O, 2 Million cells per mL). After vortexing for 30 minutes at 4°C, proteins were precipitated when centrifuging for 10 minutes at 18.000 x g at 4°C. Metabolites were resolved over the Kinetex C18 column (2.1 x 150 mm, 1.7 µm, Phenomenex) using a Vanquish UHPLC system and analyzed with the high-resolution Q Exactive mass spectrometer (Thermo Scientific) at 35 °C. A volume of 10 µl was injected for positive and negative ion mode, using a 5 minutes gradient at 450 µl/min from 5% to 95% of ACN/0.1% Formic Acid In Water/0.1% Formic Acid (positive mode) and 95% ACN/5% water/1mM ammonium acetate in 5% ACN/95% water/1mM ammonium acetate (negative mode), as previously reported^62^. To quantify CoA-metabolites, Cell pellets were resuspended in 400 uL MeOH and vortexed for 10 seconds. Next, 200 uL of ACN was added and sample was vortexed for 10 seconds. Supernatant was transferred to LC vials and dried down under vacuum, followed by reconstitution in 35 uL MeOH. The supernatant containing metabolites was resolved over an Acquity UPLC BEH (1.7um 2.1×100mm, Waters) column using a Vanquish UHPLC system and analyzed with the Orbitrap Exploris 120 mass spectrometer (Thermo Scientific) at 35 °C. A volume of 10 µl was injected for positive ion mode, using a 7 minutes gradient at 450 µl/min using buffer A (5mM ammonium acetate @ ph = 5.6) and 2% to 95% of buffer B (95% ACN, 5% H2O, 5mM ammonium acetate). The gradient was as follows: 0–0.5 min, 2% B; 0.5-4.0 min, 2–95% B; 4.0-5.0 min, 95% B; 5.0-5.2 min, 95-2% B; followed by 1.8 min re-equilibration at 2% B. An H-ESI source was utilized with positive:negative ionization at 4000:2000 V, sheath gas at 50, aux gas at 10, orbitrap resolution of 120000, and a scan range of 500-1200 m/z. Raw files were converted to mzXML file format using Raw converter and technical replicates were used to control technical variability. Only metabolites with a coefficient of variation (CV = SD/mean) < 20% were considered for this report. Metabolite assignment to KEGG compounds was performed using El-Maven, and normalization to protein content measured with the BCA protein assay kit (Thermo Scientific). Data was visualized using Metaboanalyst.

### Targeted metabolic assays

For targeted metabolic assays, induced neurons (iNs) were FACS-purified on day 21 of conversion and replated at a density of 75,000 iNs/cm². Cells were maintained either in standard neuronal conversion medium (baseline condition), or treated with 5 µM SB-204990 (MedChemExpress, Cat. No. HY-16450) starting on day 16 of conversion. Vehicle control cultures received DMSO at a final dilution of 1:2000 to match the concentration used in SB-204990-treated samples. Media and compounds were replenished at each media change until harvest. Cells were collected according to the manufacturer’s protocols for the Citrate Assay Kit (MilliporeSigma, MAK333) and the PicoProbe Acetyl-CoA Assay Kit (Fluorometric; Abcam, ab87546).

### Acetyl-CoA sensors

Genetically encoded acetyl-CoA sensors were obtained from Addgene, including mitochondrial PancACe (#215710) and mitochondrial cpGFP (#215711), nuclear PancACe (#215706) and nuclear cpGFP (#215707), as well as cytoplasmic PancACe (#215708) and cytoplasmic cpGFP (#215709)^36^. Lentiviral particles were generated by co-transfecting HEK293T cells with the respective sensor plasmids together with psPAX2 (Addgene #12260) and pMD2.G (Addgene #12259) packaging plasmids using polyethyleneimine (PEI). Viral supernatants were collected and concentrated by ultracentrifugation. FACS-purified iNs were replated on day 21 of conversion onto Geltrex-coated Culture-Insert 4 Well µ-Dish 35 mm high imaging dishes (ibidi) and transduced with the indicated sensors. Initial validation experiments assessed sensor selectivity by comparing PancACe and cpGFP control constructs following treatment with deacetylase inhibitors across control and AD conditions. As cpGFP fluorescence remained stable across conditions, subsequent experiments were performed using PancACe sensors only. Live-cell imaging was performed using an Andor Dragonfly imaging system maintained at 37 °C and 5% CO_₂_. Imaging was additionally performed on fixed samples where indicated. Cells were treated with the respective compounds for 1-48 hours prior to image acquisition and analysis. Treatments included DMSO vehicle control, ACLY inhibitor SB-204990 (5 or 10 µM), Ac-CoA Synthase Inhibitor 1 (ACSS2 inhibitor; 50 µM, Selleckchem, S8588), sodium dichloroacetate (DCA, 20 µM, Sigma, #347795), SRT1720 (Sigma-Aldrich, 5 µM, 567860), and nicotinamide riboside (NR; 200 µM, ChromaDex).

### Capillary western blot analysis

ACLY protein levels were quantified in lysates obtained from FACS-purified iNs using the ProteinSimple Jess system (Bio-Techne) with the 12-230 kDa Jess Separation Module in the chemiluminescence channel. Cell pellets were lysed in RIPA lysis and extraction buffer (Thermo Fisher Scientific) at a density of 200,000 iNs per 100 µL buffer supplemented with cOmplete EDTA-free Protease Inhibitor Cocktail (Merck, 11873580001). Samples were processed according to the manufacturer’s instructions. Primary antibodies anti-ACLY antibody (Santa Cruz Biotechnology, sc-517267, 1:50), anti-PDH-E1a (Santa Cruz Biotechnology, sc-377092, 1:50), anti-phosphorylated-PDH Ser293 (pPDH, Novus Biologicals, NB110-93479, 1:50) were incubated for 60 minutes, followed by standard default run settings provided by ProteinSimple. Data were analyzed using Compass software.

### Long-read RNA sequencing

iNs from three control donors and four AD patients were MACS purified on day 21 of conversion and subsequently replated onto Geltrex-coated 24-well plates at a density of 500,000 cells per well. The following day, medium was replaced with neuronal conversion medium supplemented with either DMSO control or ACLY inhibitor (ACLYi; 10 µM). Cells were maintained under these conditions for seven days, with three full media changes performed during treatment. Cells were then detached using TrypLE and resuspended in PBS supplemented with 0.04% BSA. For multiplexed single-cell profiling, 20,000 cells per donor line were pooled across donors per condition. Single-cell transcriptomes were captured using the 10x Genomics Chromium platform (v3.1 chemistry dual index kit) using default conditions. PacBio long-read libraries were prepared from pooled samples per condition (DMSO or ACLYi). A modified MAS-Seq workflow was used to generate full-length transcript information from 3′-barcoded cDNA libraries. Approximately 23,000 cells per sample were targeted for each condition. Sequencing libraries were generated to an average depth of ∼80 million segmented reads and long-read sequencing was performed on the PacBio Revio system. Genomic DNA was extracted from wild-type fibroblasts using the Qiagen DNeasy Blood & Tissue Kit, and DNA concentration and quality were assessed using a NanoDrop spectrophotometer. Single nucleotide polymorphism (SNP) profiles were assessed using the Infinium CoreExome-24 (Illumina) and used to demultiplex pooled single-cell datasets and assign cells to their respective donors using cellsnp-lite coupled with vireo^63,64^. Dimensionality reduction and subsequent clustering was generated using Seurat^65^. For UMAP, top 30 PCs were used for initial clustering. Identification and filtering of excitatory and inhibitory neurons was based on well-established maker genes. Subsequently, filtered cells were re-clustered. Cell-type-specific and filtered barcodes were extracted from the Seurat-object to generate subtype-resolved BAM files for each donor and condition combination, enabling downstream isoform-level analysis in a cell-type-restricted pseudobulk manner. To ensure sufficient sequencing coverage for reliable quantification, clusters containing fewer than 100 cells were excluded from downstream analysis. Raw BAM files were processed by removing non-standard and unplaced contigs and subsequently sorted and indexed using SAMtools. Reads were aligned to the GRCh38.p13 reference genome using ESPRESSO_S^32^, with the GENCODE v39 basic annotation as reference. Splice junction correction and transcript model refinement were performed using ESPRESSO_C to recover and refine full-length isoforms and novel splice junctions across samples. Transcript-level quantification outputs from ESPRESSO were used for downstream differential expression and isoform usage analyses. Differential isoform expression was assessed using DESeq2 in R^66^, and multiple testing correction was performed using the Benjamini-Hochberg procedure within each gene. For alternative splicing and isoform structure comparisons, isoforms were identified across all samples simultaneously, including excitatory and inhibitory neurons from both DMSO and ACLYi conditions. Resulting event-level outputs (.esp files) were harmonized in R to ensure consistent transcript/isoform identifiers across conditions, including novel ESPRESSO-defined isoforms. Alternative splice events were identified using rMATS_long^31^. Differential isoform usage and expression changes were evaluated using Fisher’s exact test and model-based frameworks depending on output type. Rescue effects under ACLYi treatment were defined as a reversal of the direction of log2 fold-change between DMSO and ACLYi conditions relative to baseline comparisons. All analyses were performed at transcript resolution using long-read transcript identifiers to preserve isoform specificity.

### Acetyl-proteomics

Acetyl-proteomics was performed across two independent experimental runs. The initial dataset comprised 11 control and 7 Alzheimer’s disease (AD) samples and was analyzed using spectral count-based quantification. The second dataset included 7 control and 6 AD samples, each measured at baseline and following treatment with either DMSO or the ATP citrate lyase inhibitor (ACLYi; 10 µM, 7 days). This dataset was quantified using MAX-LFQ-normalized intensity values. One million iNs were FACS/MACS purified, snap-frozen. Acetylated peptides were enriched using the PTMScan® HS Acetyl-Lysine (Ac-K) Kit (Cell Signaling Technology, Danvers, MA) according to the manufacturer’s protocol with minor modifications. Briefly, protein lysates were prepared in urea lysis buffer containing protease and deacetylase inhibitors, reduced with dithiothreitol (DTT), alkylated with iodoacetamide (IAA), and digested overnight with trypsin. The resulting peptides were desalted and dried by vacuum centrifugation. Peptides were resuspended in HS IAP bind buffer and incubated with the immobilized anti–acetyl-lysine antibody beads at 4°C for 2 hours on an end-over-end rotator. Following enrichment, the beads were washed sequentially with HS IAP wash buffer and water to remove nonspecific binders. Bound peptides were eluted with 0.15% trifluoroacetic acid (TFA), desalted using C18 StageTips, and dried by vacuum centrifugation prior to LC–MS/MS analysis. The samples were analyzed on a nano-UHPLC (nanoElute, Bruker Daltonics) coupled to a timsTOF SCP mass spectrometer (Bruker Daltonics). Samples were separated on an Aurora Ultimate column 25 cm x 75 mm, C18 1.7 mm (IonOptics) which was heated to 50 °C in a column oven. Mobile phases consisted of 0.1% (v/v) formic acid in water (phase A) and acetonitrile (phase B). Samples were separated on a 90 min stepped gradient ranging from 4-35% B at a flow rate of 300 nL/min. The gradient is built of sequential steps ranging from 4 to 25% B in 60 min, 25 to 35% B in 20 min, and finally, 35 to 95% in 10 min. The performance of the mass spectrometer was monitored by standardized measurements of a K562 tryptic digest (Pierce), where the number of peptides and the total ion chromatogram (TIC) shape were considered. The mass spectrometer was operated in PASEF mode. The accumulation and ramp time was set to 166 ms and 10 PASEF MS/MS scans per topN acquisition cycle were acquired. MS and MS/MS spectra were recorded from m/z 100 to 1700. The ion mobility was scanned from 0.6 to 1.40 Vs/cm2. Precursors for data-dependent acquisition were isolated within ± 1 Th and fragmented with an ion mobility-dependent collision energy, which was linearly increased from 20 to 65 eV in positive mode. Low-abundance precursor ions with an intensity above a threshold of 1000 counts but below a target value of 12500 counts were repeatedly scheduled and otherwise dynamically excluded for 0.2 min.

Raw data files conversion to peak lists in the MGF format, downstream identification, validation, filtering and quantification were managed using FragPipe version 13.0. MSFragger version 3.0 was used for database searches against a human database with decoys and common contaminants added. Search parameters were set for trypsin-specific cleavage allowing up to two missed cleavages, with up to two isotope errors permitted in precursor selection. The precursor and fragment mass tolerances were set to 25 ppm (MS1) and 0.04 Da (MS2), respectively. Dynamic modifications included oxidation of methionine (+15.9949 Da), Dynamic modifications included oxidation of methionine and proline (+15.9949 Da), lysine acetylation (+42.0106 Da), N-terminal acetylation (+42.0106 Da), carboxymethylation (+57.0215 Da), and lysine ¹³C_₂_ acetylation (+44.0173 Da).

All datasets were log-transformed prior to downstream analysis. Proteins with zero intensity across all samples were excluded. In addition, samples exhibiting high intra-group variability (standard deviation > 30 within group) were removed to reduce technical noise and improve robustness of downstream inference. For global acetylation assessment, summed signal intensities across all detected acetylated proteins were calculated per sample. Additionally, the total signal of acetylated proteins was normalized to the corresponding non-acetylated protein fraction (supernatant of the unbound fraction) to confirm that increased acetylation levels were not driven by technical artifacts arising from differences in overall protein abundance in the AD group. Because not all proteins detected in the acetylated fraction were also detected in the non-acetylated fraction, a per-protein normalization strategy would have excluded a substantial proportion of proteins exhibiting disease-associated changes. Therefore, we focused on the acetyl-enriched fraction in order to avoid introducing bias and potentially discarding biologically relevant alterations associated with disease state. Differential acetylation analysis was performed using linear modeling as implemented in the limma R package. Log2-transformed acetylation intensities were modeled using a design matrix without intercept, with each experimental condition represented as an individual coefficient. To account for donor-matched repeated measurements, donor identity was included as a blocking factor, and within-donor correlation was estimated using the duplicateCorrelation function. The resulting correlation structure was incorporated into the linear model fitting procedure. Specific contrasts were defined to assess: (i) disease-associated differences (AD vs control), (ii) treatment effects within AD (ACLYi-treated vs AD baseline), (iii) treatment effects within controls (ACLYi-treated vs control baseline), and (iv) cross-condition treatment comparisons (AD-treated vs control-treated). Empirical Bayes moderation was applied to improve variance estimation and statistical stability. Subcellular localization of differentially acetylated proteins was annotated using the Human Protein Atlas subcellular localization database (proteinatlas.org, accessed April 2025).

### Total Proteomics

Purified induced neurons (iNs) were lysed in N-PER extraction buffer (Thermo Scientific) according to the manufacturer’s instructions. Protein concentration was determined by BCA assay, and equal amounts of protein were used for downstream processing (approximately 30 µg per sample). Disulfide bonds were reduced using tris(2-carboxyethyl)phosphine (TCEP), followed by alkylation with 2-chloroacetamide (CAA). Proteins were digested overnight with trypsin at 37°C. Resulting peptides were desalted and subjected to tandem mass tag (TMT) labeling for multiplexed quantitative proteomics. Following labeling, samples were pooled and fractionated by high-pH reversed-phase liquid chromatography prior to mass spectrometry analysis. Peptides were analyzed on an Orbitrap Eclipse mass spectrometer coupled to an Easy-nLC 1200 system (Thermo Scientific). Raw data were acquired in data-dependent acquisition mode and processed for quantitative comparison across conditions. Protein abundance was normalized based on total peptide input, with additional verification using BCA-derived loading equivalence across samples. After removal of BSA and Albumin counts, the samples were median normalized and log-transformed before calculating differential protein levels using limma.

## Statistical analysis

Normality of data was assessed using the Shapiro-Wilk or Kolmogorov-Smirnov tests. Two-tailed statistical tests were used unless otherwise stated. Bar plots display mean ± standard deviation; violin plots show the median and interquartile range. If sample size was greater than 1 and less than 100, individual data points are shown as dots. Statistical significance is indicated as follows: *p* < 0.05 (*)*, p < 0.01 (**), p < 0.001 (\*\*\**), and *p* < 0.0001 (****). ANOVA and *t*-tests were performed using GraphPad Prism. Linear regression and correlation analyses were performed using GraphPad Prism and RStudio.

