## Supplementary material for "Nuclear acetyl-CoA accumulation drives reversible RBP hyperacetylation and transcriptome-wide splicing shifts in age-equivalent Alzheimer’s disease neurons": Document S1

**SUPPLEMENTARY FIGURES**

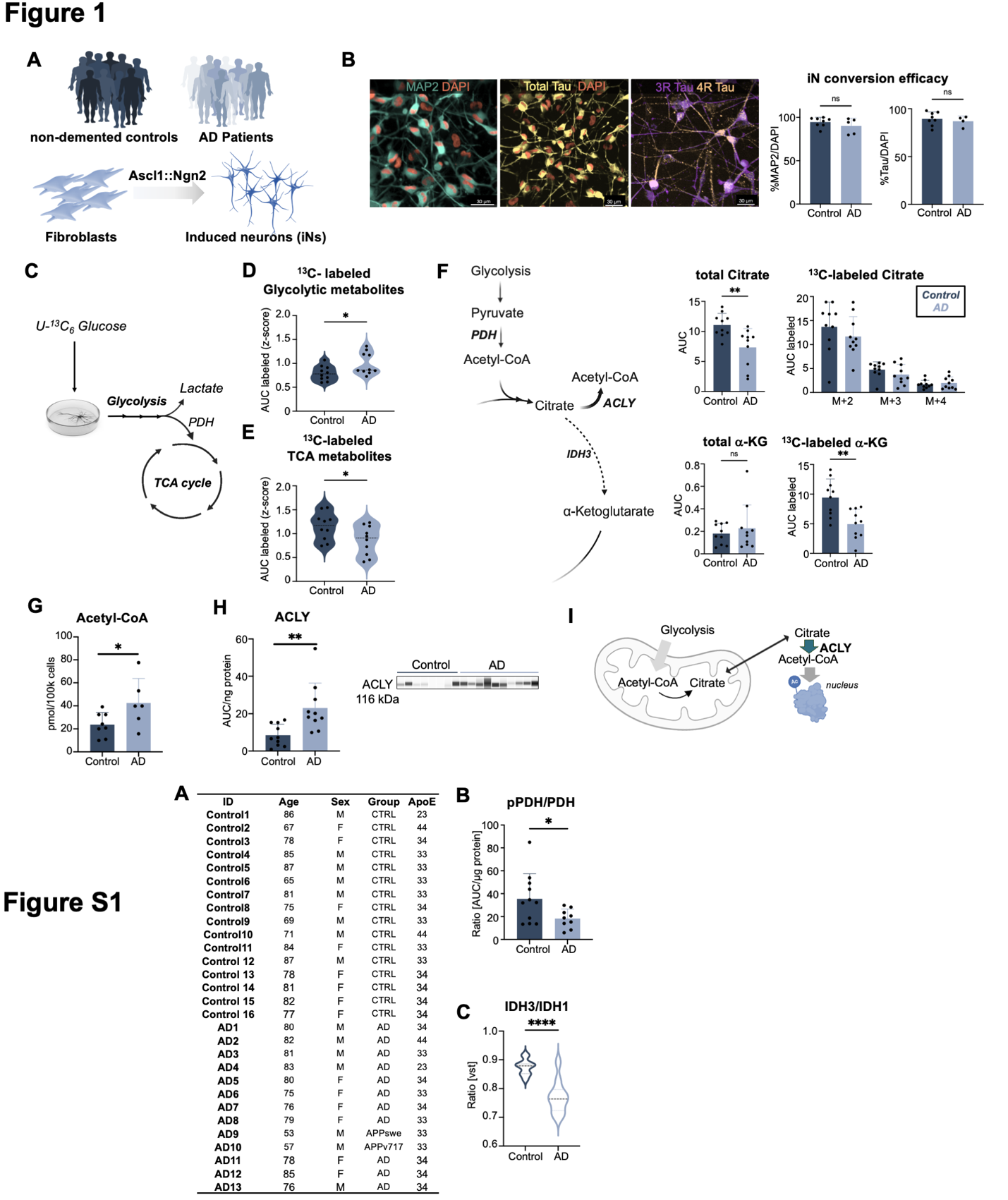

**Figure S1. Metabolic re-wiring in AD iNs.**

(A) Cohort overview.

(B) ProteinSimple capillary western blot analysis of pyruvate dehydrogenase (PDH) and phosphorylated PDH (pPDH, Ser293) in directly converted induced neurons (iNs). Signal intensities were normalized to total protein measured by BCA assay. n = 11 control donors and n = 9 AD donors. Each dot represents an independent donor. Data are presented as mean ± SD.

(C) RNA-seq analysis of a previously published dataset^15^. Raw counts were normalized using variance-stabilizing transformation (VST), and IDH3-to-IDH1 expression ratios were calculated. n = 11 control donors and n = 10 AD donors. Black dotted line indicates the median; blue dotted lines indicate quartiles.

(B,C) Statistical significance was determined using an unpaired two-tailed Student’s t-test.

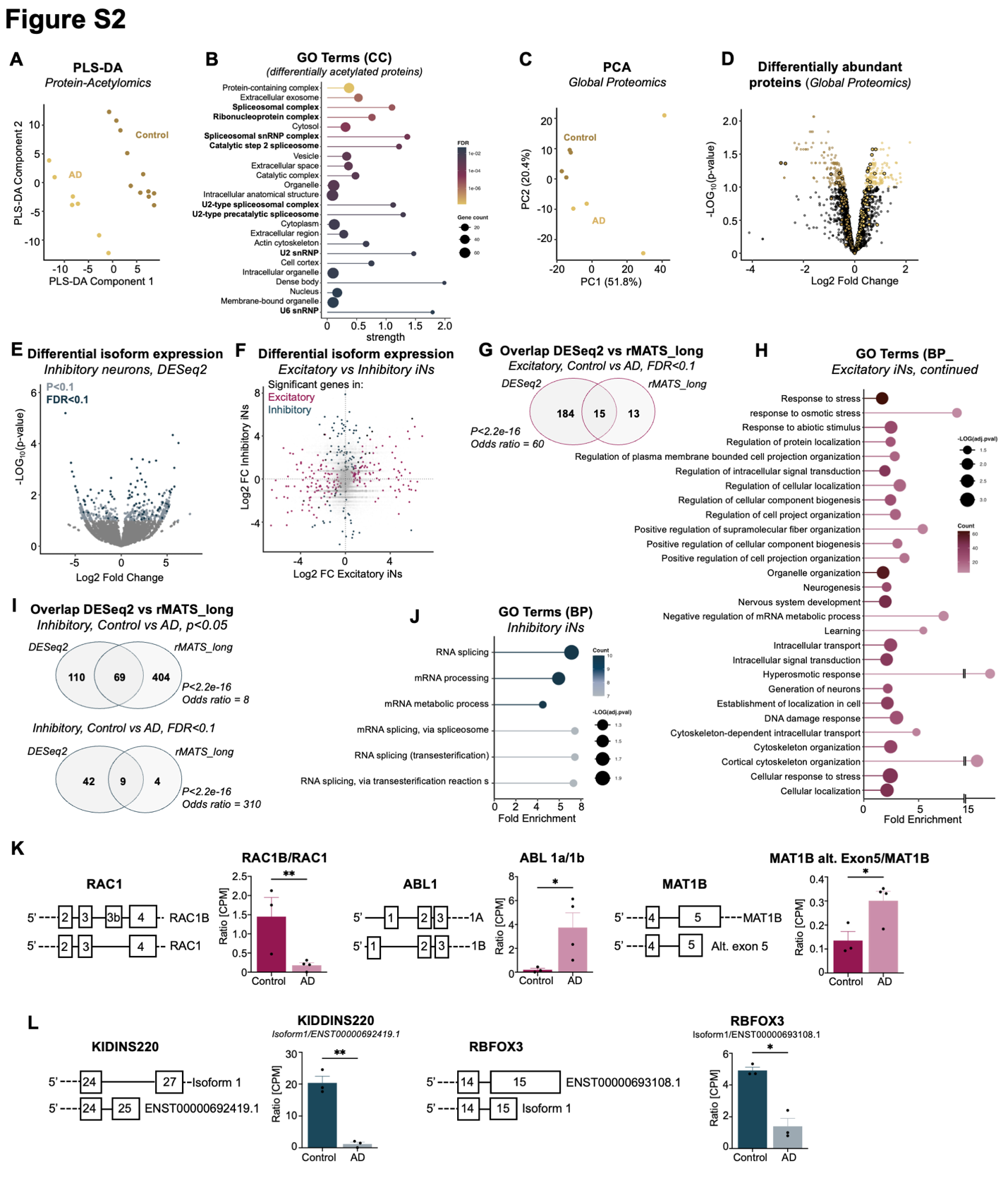

**Figure S2. Hyperacetylation of RBPs and alternative splicing in AD iNs.**

(A) PLS-DA clustering of global protein acetylomics. Acetylated proteins were enriched using the PTMScan Acetyl-Lysine Motif Kit and analyzed by mass spectrometry. n = 11 control donors and n = 7 AD donors.

(B) Differential acetylation analysis performed using limma on log-transformed data. Gene Ontology cellular compartment (GO-CC) enrichment analysis was performed using STRING. Shown are all significant terms with a false discovery rate (FDR) < 0.05.

(C) PCA clustering of TMT-based quantitative proteomics. n = 4 control donors and n = 4 AD donors.

(D) Differential protein abundance analysis using limma on median-normalized and log-transformed data comparing control versus AD iNs. RBPs are highlighted with black circles.

(E) DESeq2 differential gene expression analysis in inhibitory neurons showing all detected transcripts. Transcripts with p < 0.1 are highlighted in light blue and genes with FDR < 0.1 in dark blue. Comparison shown is control versus AD iNs. n = 3 control donors and n = 3 AD donors.

(F) Correlation of log2 fold changes (log2FC) in excitatory versus inhibitory neurons comparing control and AD iNs. Transcripts significant in excitatory neurons (FDR < 0.1) are highlighted in red, and transcripts significant in inhibitory neurons (FDR < 0.1) are highlighted in blue.

(G) Overlap of significantly altered transcripts identified by DESeq2 and rMATS-long analysis in excitatory neurons. All transcripts passing an FDR threshold < 0.1 are shown. Statistical significance of overlap was calculated using Fisher’s exact test.

(H) DAVID functional annotation analysis of overlapping genes that are alternatively spliced in AD iNs identified by DESeq2 and rMATS-long analysis, extended from Figure 2K.

(I) Overlap of significantly altered transcripts identified by DESeq2 and rMATS-long analysis in inhibitory neurons. Shown are all transcripts passing either a nominal p < 0.05 threshold (top panel) or an FDR < 0.1 threshold (bottom panel). Statistical significance of overlap was calculated using Fisher’s exact test.

(J) DAVID functional annotation analysis of overlapping genes that are alternatively spliced in AD iNs identified by DESeq2 and rMATS-long analysis. Shown are all significant Gene Ontology biological process (GO-BP) terms. No significant WikiPathways terms were identified.

(K) Representative genes differentially spliced in AD iNs in excitatory neurons. Statistical significance was determined using unpaired two-tailed t-tests.

(L) Representative genes differentially spliced in AD iNs in inhibitory neurons. Statistical significance was determined using unpaired two-tailed t-tests.

(K,L) Each dot represents an independent donor. Data are presented as mean ± SD.

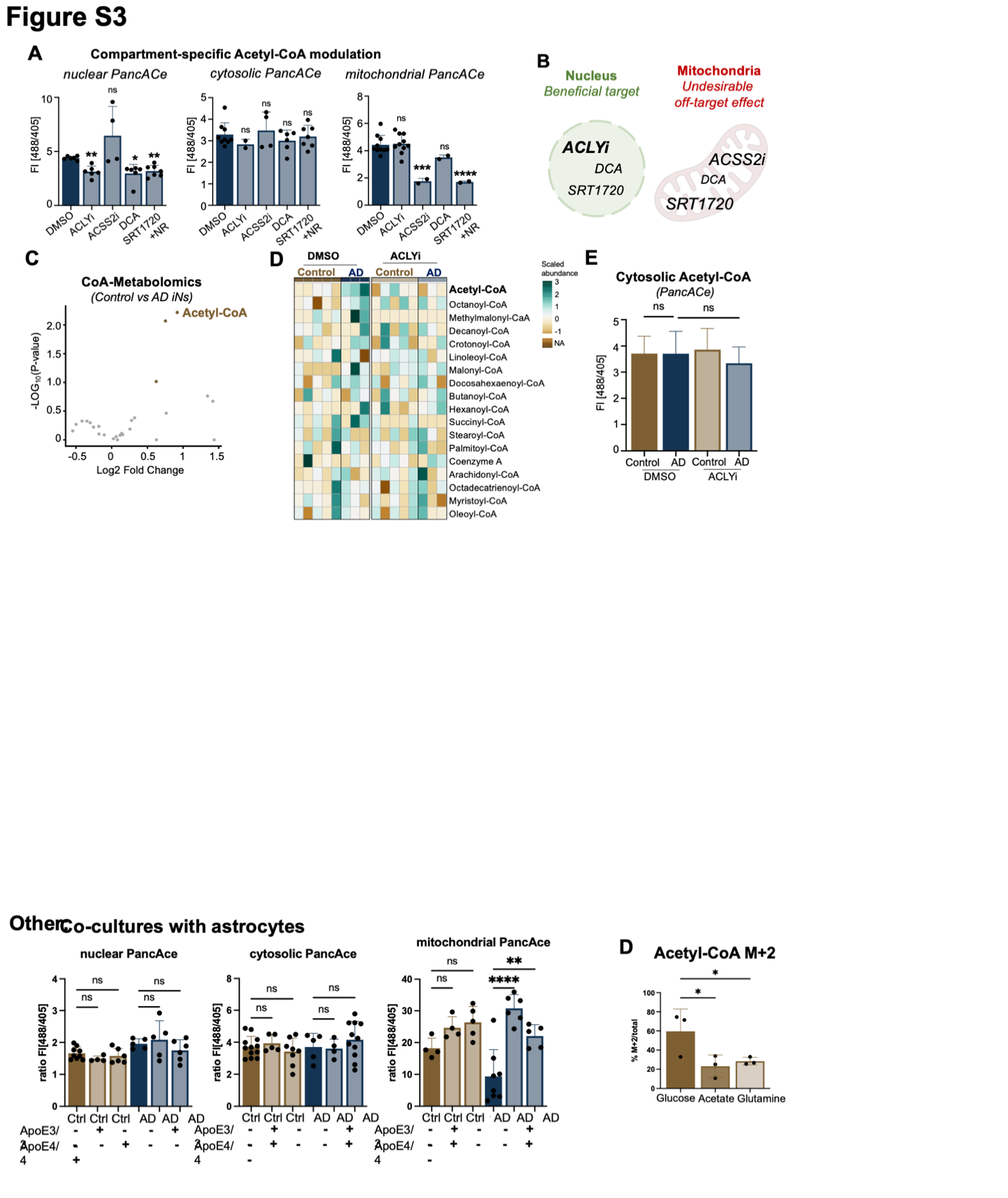

**Figure S3. Inhibition of ACLY restores subcellular acetyl-CoA homeostasis and mitochondrial metabolism in AD iNs.**

(A) Compound screen assessing the effects of metabolic inhibitors on compartmentalized acetyl-CoA sensor signals in fixed cells. n = 2-10 donors per condition; each dot represents an independent donor. Statistical significance was determined using ordinary one-way ANOVA with Dunnett’s multiple comparisons test (cytosolic: F(4,23) = 0.67; mitochondrial: F(4,21) = 15.0) or Brown-Forsythe and Welch ANOVA tests with Dunnett’s T3 multiple comparisons test (nuclear PancACe did not pass Shapiro-Wilk and Kolmogorov-Smirnov normality tests; Brown-Forsythe ANOVA: F(4,4) = 5.4; Welch’s ANOVA: W(4,9.7) = 14.9).

(B) Schematic summarizing the effects of different metabolic compounds on compartmentalized acetyl-CoA pools.

(C) UHPLC-MS quantification of CoA metabolites under baseline conditions (DMSO). n = 5 control donors and n = 3 AD donors.

(D) Heatmap showing z-scores of all detected CoA metabolites quantified by UHPLC-MS.

(E) Quantification of cytosolic PancACe sensors in control and AD iNs under baseline conditions (DMSO) or following 7 days of ACLYi treatment. Sensor intensity was measured using 488 nm excitation and normalized to the internal control signal excited at 405 nm. n = 12 control DMSO, n = 5 AD DMSO, n = 4 control ACLYi, and n = 7 AD ACLYi donors. Statistical significance was determined using ordinary one-way ANOVA (F(3,24) = 0.6) followed by Sidak’s multiple comparisons test. Scale bar = 20 µm.

Data are presented as mean ± SD.

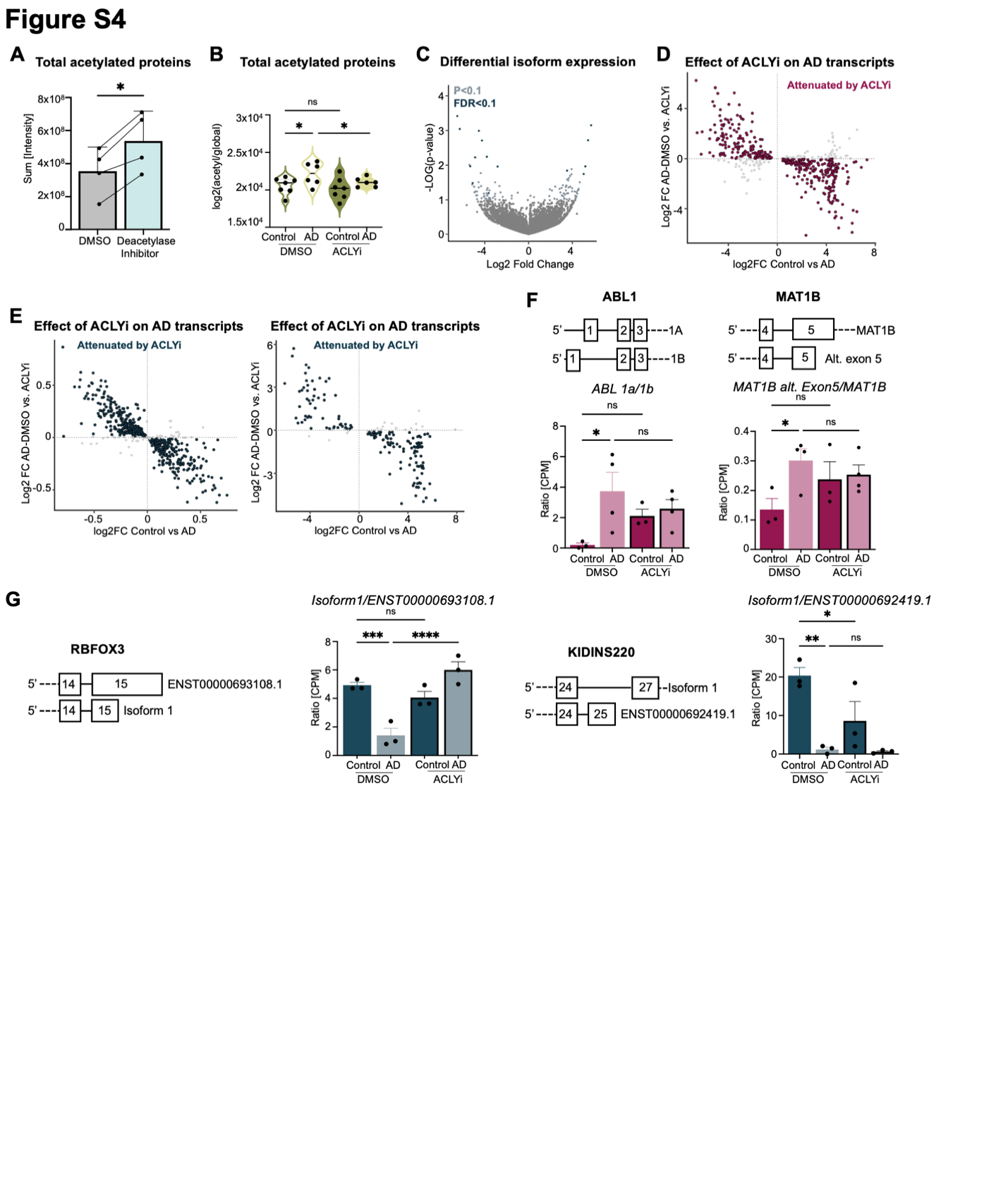

**Figure S4. ACLY inhibition reverses hyperacetylation of RBPs and restores a healthy RNA isoform landscape.**

(A) Acetylated proteins were enriched using the PTMScan Acetyl-Lysine Motif Kit and analyzed by mass spectrometry. iNs were MACS-purified and treated with either DMSO or a deacetylase inhibitor mix for 24 h. Shown is the total number of acetylated proteins detected per condition. n = 4 DMSO-treated iN cultures and n = 4 matched deacetylase inhibitor-treated cultures. Statistical significance was determined using paired two-tailed t-tests. Matched donors are indicated by connecting lines.

(B) Total acetylated proteins normalized to the total number of proteins identified in the flow-through (global proteome) fraction following enrichment of acetylated proteins. n = 7 control donors (DMSO/ACLYi) and n = 6 AD donors (DMSO/ACLYi). Statistical significance was determined using ordinary one-way ANOVA (F(3,22) = 3.1). Paired two-tailed t-tests were additionally performed to compare matched AD DMSO and AD ACLYi samples, indicated by connecting lines.

(C) DESeq2 differential transcript expression analysis in inhibitory neurons comparing AD DMSO versus AD ACLYi conditions. All detected transcripts are shown; genes with p < 0.1 are highlighted in light blue and genes with FDR < 0.1 in dark blue.

(D) Correlation of isoform changes between control versus AD (DMSO; x-axis) and AD DMSO versus AD ACLYi (y-axis) in excitatory neurons. Each dot represents a transcript significantly altered (FDR < 0.1) in the control versus AD DMSO comparison. Red dots indicate transcripts shifted in the opposite direction following ACLYi treatment (rescue).

(E) Correlation of isoform changes between control versus AD (DMSO; x-axis) and AD DMSO versus AD ACLYi (y-axis) in inhibitory neurons. Each dot represents a transcript significantly altered in the control versus AD DMSO comparison (left: p < 0.05; right: FDR < 0.1). Blue dots indicate transcripts shifted in the opposite direction following ACLYi treatment (rescue).

(F) Differentially spliced in AD iNs in excitatory neurons. Statistical significance was determined using ordinary one-way ANOVA followed by paired two-tailed t-tests comparing matched AD DMSO and AD ACLYi samples. ABL1: F(3,10) = 3.1; MAT1B: F(3,10) = 2.6.

(G) Differentially spliced in AD iNs in inhibitory neurons. Statistical significance was determined using ordinary one-way ANOVA followed by paired two-tailed t-tests comparing matched AD DMSO and AD ACLYi samples. KIDDINS220: F(3,8) = 11.3; RBFOX3: F(3,8)=18.8.

Each dot represents an independent donor. Data are presented as mean ± SD. Violin plots show the median (black line) and quartiles (colored dotted lines).
